# High-depth whole genome sequencing of blood culture plates reveals evolutionary dynamics in cases of persistent bacteremia due to methicillin-resistant *Staphylococcus aureus*

**DOI:** 10.64898/2026.04.25.720776

**Authors:** Emma G. Mills, Leah M. Grady, Edwin Chen, Marla G. Shaffer, Ryan K. Shields, Daria Van Tyne, Matthew J. Culyba

**Author notes:** Address correspondence to Matthew J. Culyba. Department of Microbiology and Immunology, Roy J. and Lucille A. Carver College of Medicine, University of Iowa, Iowa City, IA.

## Abstract

Within a single bacterial strain, DNA sequence variation is expected between individual clones. Whole genome sequencing (WGS) can be applied to clinical cultures to detect this polyclonal variation, enabling tracking of within-host evolution and transmission. Culture isolates from infected patients are often sequenced as individual colonies (c-seq). To increase the sensitivity of variant detection, cultures can also be sequenced to a high depth of coverage as a pool (p-seq), but the utility of this approach is not clear for most clinical specimens. To understand the performance of high-depth WGS in bacteremia, we applied p-seq to blood culture plates for 10 patients with persistent bacteremia due to methicillin-resistant *Staphylococcus aureus*. As a comparison, for six patients, we also applied c-seq to five colonies (c5-seq) from the same plates. p-seq was more sensitive than c5-seq for detecting low frequency variant alleles; however, the most important factor for new variant detection was the number of culture plates analyzed rather than the sequencing method used. We also used these data to construct Muller plots for three patients with especially diverse infecting populations, which enabled visualization of rapid evolutionary dynamics in response to antibiotic exposures. We identified 204 unique variant alleles, and our analysis provides additional evidence for parallel evolution of several different genes during *S. aureus* bacteremia. Overall, these data provide a detailed view of evolutionary dynamics during clinical cases of MRSA bacteremia and describe the merits and limitations of a c-seq versus p-seq strategy for analyzing blood culture plates using WGS.

**Importance:** As bacterial whole genome sequencing (WGS) is increasingly used as a research tool for clinical samples, it is important to understand the pros and cons of different culture sampling methodologies. Here, we analyzed cases of persistent bacteremia due to methicillin-resistant *Staphylococcus aureus* by applying WGS to either each of five individual colonies isolated on blood culture plates (c5-seq) or the pooled bacterial population on each plate (p-seq). We found that c5-seq was a more practical and informative method to understand evolutionary dynamics.

## Introduction

The application of whole genome sequencing (WGS) to clinical bacterial isolates continues to expand our understanding of bacterial adaptation during human infection and colonization (1). It is important to optimize the approach to sequencing. Even within a single bacterial strain, DNA sequence variation is expected between clones. WGS can be used to characterize this ‘polyclonal’ variation, enabling investigators to infer transmission events between patients or to track the within-host evolution of a pathogen through time. In such studies, differences of <100 single nucleotide polymorphisms (SNPs) typically describe the polyclonal variation within a single infecting or colonizing strain (2, 3). The standard approach to characterize this variation is to pick individual colonies from culture plates, sequence the genome of each colony separately, then compare their genome sequences. We refer to this method as colony-seq (c-seq). With c-seq, the sensitivity of allele detection is a function of the number of colonies sequenced (**Fig. 1A**). For blood cultures, to balance sensitivity with cost, sequencing five individual colonies (c5-seq) has been proposed (4). However, when sampling a culture plate with a large population size, statistically, c5-seq should only reliably (≥95% confidence) detect variant alleles that are present on the plate at an allele frequency (AF) of ≥50%. Alternatively, the plate population can be sequenced to a high depth of coverage as a pool (p-seq) (**Fig. 1B**). In this approach, since bacterial population sizes on blood culture plates are large, sensitivity is no longer limited by the number of colonies sequenced from the plate, but instead by the inherent base-calling error rate of next generation sequencing (e.g. ∼1% for Illumina platforms) (5) and errors associated with read alignment to the reference genome. With careful filtering of false-positive variant calls, this approach is reported to detect variants with an AF as low as 3% (6), which is significantly more sensitive than c5-seq. However, for clinical blood cultures, the blood sample is first cultured in liquid broth prior to streak-plating on solid media (i.e. blood culture plate) to obtain isolated colonies (**Fig. 1B**). Importantly, initial blood inoculum sizes are estimated to vary over a wide range (1 to 1000 CFU) (7). Inoculum sizes on the low end of this range could place such a severe sampling bottleneck on variant allele detection that the added sensitivity of p-seq is rendered entirely moot, particularly if the genetic diversity within the infecting bacterial population is low. Thus, the added utility of employing p-seq for blood culture plates is not clear.

**Figure 1.**
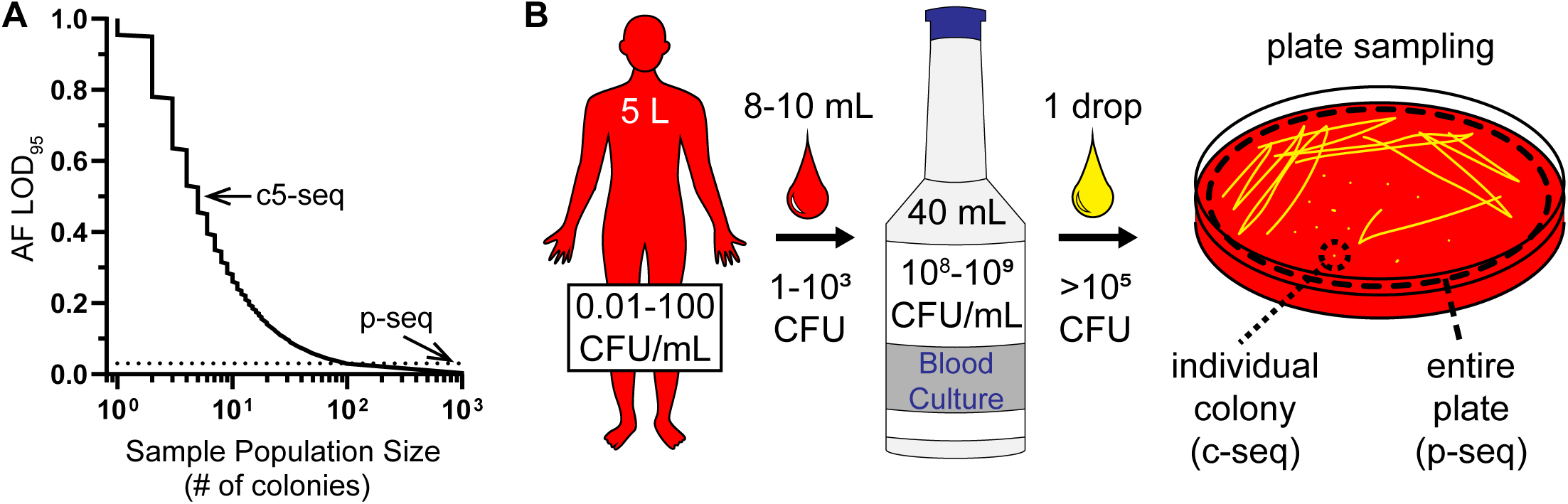
Allele detection sensitivity and sampling bottlenecks of WGS on blood culture plates. **A.** The plot shows the lowest allele frequency that can be detected with 95% confidence (AF LOD_95_) as a function of the number of colonies sampled, as given by the binomial distribution. Sequencing five colonies (c5-seq) has an AF LOD_95_ = 0.5, whereas when sequencing more than ∼100 colonies (via p-seq) detection is limited by the inherent variant-calling errors of WGS (AF ∼ 0.03, dotted horizontal line). **B.** For clinical blood cultures, the initial inoculum size of the blood sample (8-10 mL) is relatively small (1-10^3^ CFU) compared to an adult patient’s blood volume (5 L); and is first amplified by growth within a blood culture bottle prior to streak-plating. Individual colonies can be picked for sequencing (c-seq) or all the growth on the plate can be collected and sequenced as a pool (p-seq).

Here, we examined the performance of p-seq applied to blood culture plates for ten cases of persistent bacteremia due to methicillin-resistant *S. aureus* (MRSA). In six cases, we also applied c5-seq to the same blood culture plates to allow for direct comparison. To our knowledge, this study represents the deepest sampling of blood culture plates for individual cases of persistent bacteremia to date. We found that the number of genotypes detected varied considerably between patients, and we identified three patients with especially diverse infecting populations. We analyzed two independent blood culture plates for most time points, yielding ten individual colonies and two pooled mixtures for these time points. This enabled construction of Muller plots to visualize genotype frequency as a function of time. Major shifts in genotype frequencies correlated with antibiotic exposures, sometimes selecting for relatively more or less antibiotic-resistant strains. p-seq was overall a more sensitive method for detecting variant alleles, but the genetic linkage information from c-seq allowed for accurate characterization of genotype dynamics. Moreover, we found it was important to sample all available blood culture plates to characterize the genetic diversity of the infecting population.

## RESULTS

### Measurement performance and variant call optimization of p-seq

Before applying p-seq to clinical blood culture plates, we first determined the accuracy and precision of its AF measurements. To do this, we used two clones of *S. aureus* that differed by five SNPs and made control samples by mixing the genomic DNA (gDNA) of the two clones at different DNA ratios to achieve five expected frequencies (AF_exp_: 0.5, 0.2, 0.1, 0.05, and 0.01). Then, we sequenced each sample mixture at high coverage (mean 1073X) and ran the *breseq* variant-caller program (8) in ‘polymorphism mode’ to obtain an observed frequency (AF_obs_) for each SNP. We treated the five SNPs as replicate measurements since we expected them to have the same values of AF_obs_ within each sample mixture. The mean ± standard deviation of the AF_obs_ values for samples with AF_exp_ of 0.5, 0.2, 0.1, 0.05, and 0.01 were 0.509 ± 0.032, 0.205 ± 0.018, 0.084 ± 0.013, 0.056 ± 0.022, and 0.008 ± 0.006, respectively (**Fig. 2A**). Linear regression showed high correlation (R^2^=0.99) between AF_obs_ and AF_exp_ values. Overall, we found the error associated with p-seq AF measurements to be minimal. The mean absolute difference of all observed measurements from their expected values was 0.016 and the combined standard deviation was 0.020.

**Figure 2.**
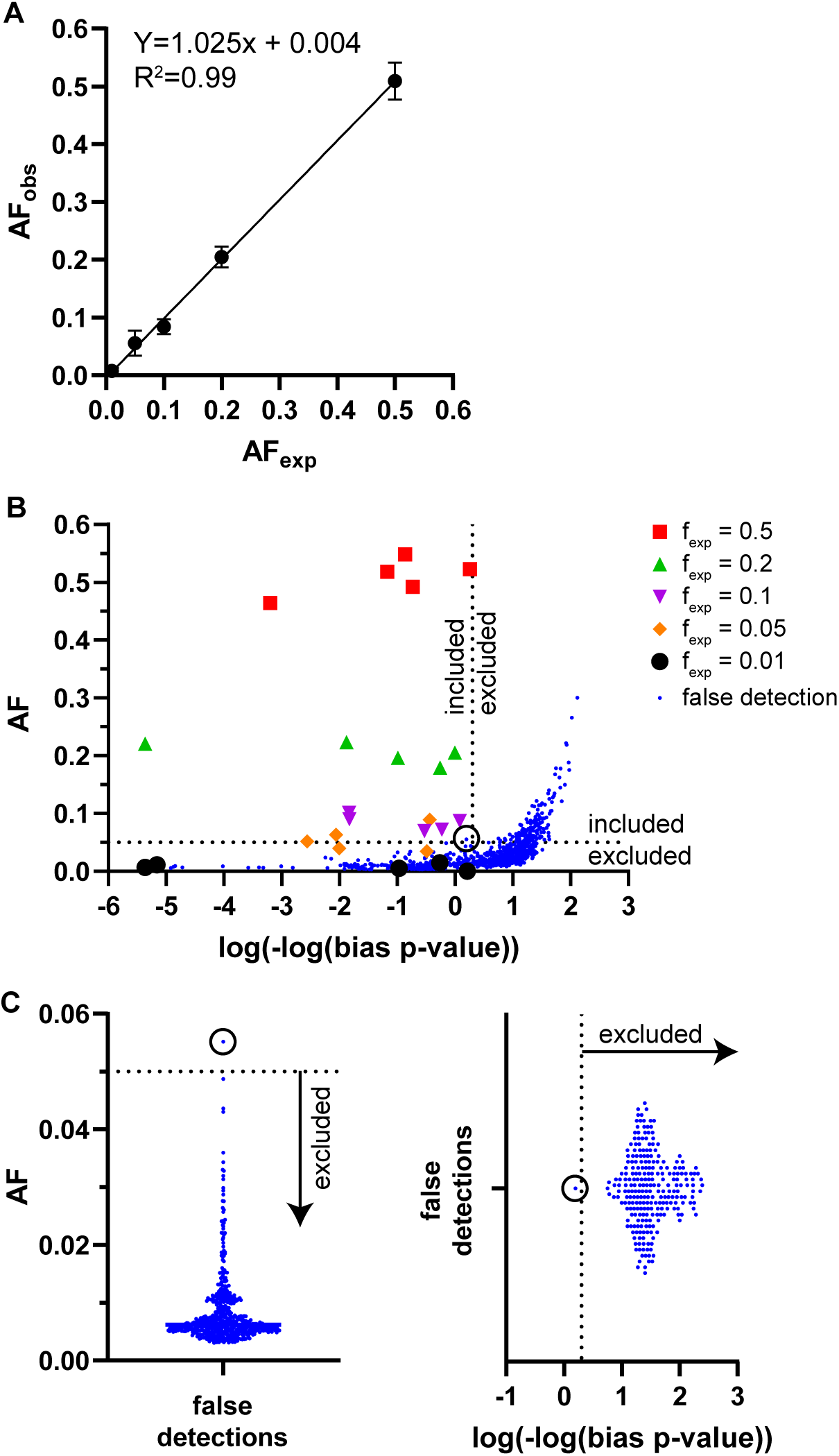
Measurement performance and variant call optimization of p-seq. **A.** Plot of observed allele frequency values (AF_obs_) versus expected allele frequency values (AF_exp_). Data points and error bars represent means and standard deviations, respectively (n=5). The line represents the best fit of linear regression. **B.** Scatter plot of AF versus log(-log(bias p-value)) for all the variant calls. Large data symbols indicate true variants and blue dots indicate false detections. Dotted lines represent threshold values chosen for variant call filtering. Only one false-positive variant is included (circled datapoint) using this filtering strategy. ‘Bias p-value’ is a *breseq* output parameter that quantifies variant-call quality. **C.** AF (left) and bias p-value (right) false detection distributions shown as column scatter plots with adjusted axis scales to more clearly show the threshold values used for variant call filtering. Data points for false detections are the same as in panel B. The plots show that only one false-positive variant is included (circled datapoint – same datapoint as in panel B) using this filtering strategy. For clarity, the plots are oriented to align with the axis orientation of panel B and datapoints excluded by both filters are not shown.

We also used the same *breseq* data output to optimize identification and filtering of false-positive variant calls, since all true-positive variants in the control samples were known. The *breseq* output parameters ‘frequency’ and ‘bias p-value’, which are associated with each variant call, were used for this purpose. The frequency parameter (referred to above as AF_obs_) is the estimate for the non-reference (variant) AF in the population. Bias p-value represents a statistical evaluation of the AF estimate, which includes the bases observed in the read pileup and their quality scores. We examined the scatter plot of AF versus log(-log(bias p-value)) for all variant calls to identify threshold values of these parameters that discriminated between true-positive and false-positive calls; and chose to exclude p-seq variant calls with AF_obs_ <0.05 and bias p-value <0.01 (**Fig. 2B**). These criteria resulted in the inclusion of only one false-positive variant call that represented the tail-end of both the AF and bias p-value distributions (**Fig. 2C**). This control dataset was comprised of five separate p-seq samples, which yields an estimate for the false-detection rate of 0.2 per sample.

### Sensitivity of variant allele detection and estimation of false-detection rate

We next applied p-seq to 109 blood culture plates derived from ten patients with persistent bacteremia due to MRSA (**Table S1**). For six patients (75 plates), we also applied c5-seq, thereby sequencing 375 individual clones in total. After analyzing WGS reads for evidence of contaminating bacteria using *kraken* (9), we excluded three samples from further analysis, yielding 107 plate samples and 374 individual clones. For each patient, a separate reference genome was assembled *de novo* using gDNA from an individual colony. To detect genetic variants, we mapped the sequencing reads from each sample to the reference genome derived from the same patient infection, thereby characterizing the within-host genetic diversity of each infection episode. As expected, p-seq was more sensitive for variant allele detection than c5-seq. Out of a total of 204 variants across the entire dataset, p-seq detected 189 (93%) and c5-seq detected 173 (85%) (**Fig. 3A**), yielding p-seq 1.1-fold (189/173) more variants than c5-seq. Notably, the number of additional unique variants detected by p-seq varied significantly among the patients. For example, p-seq detected no additional variants beyond that of c5-seq for Patients C and G, whereas it identified 11 and 13 additional alleles for Patients B and D, respectively. We also analyzed the data on a per plate basis, allowing variants to be counted multiple times if they were on more than one plate (**Fig. 3B**). This resulted in a similar distribution. Out of a total of 1,080 variants, p-seq detected 985 (91%) and c5-seq detected 886 (82%), yielding p-seq 1.1-fold (985/886) more variants than c5-seq.

**Figure 3.**
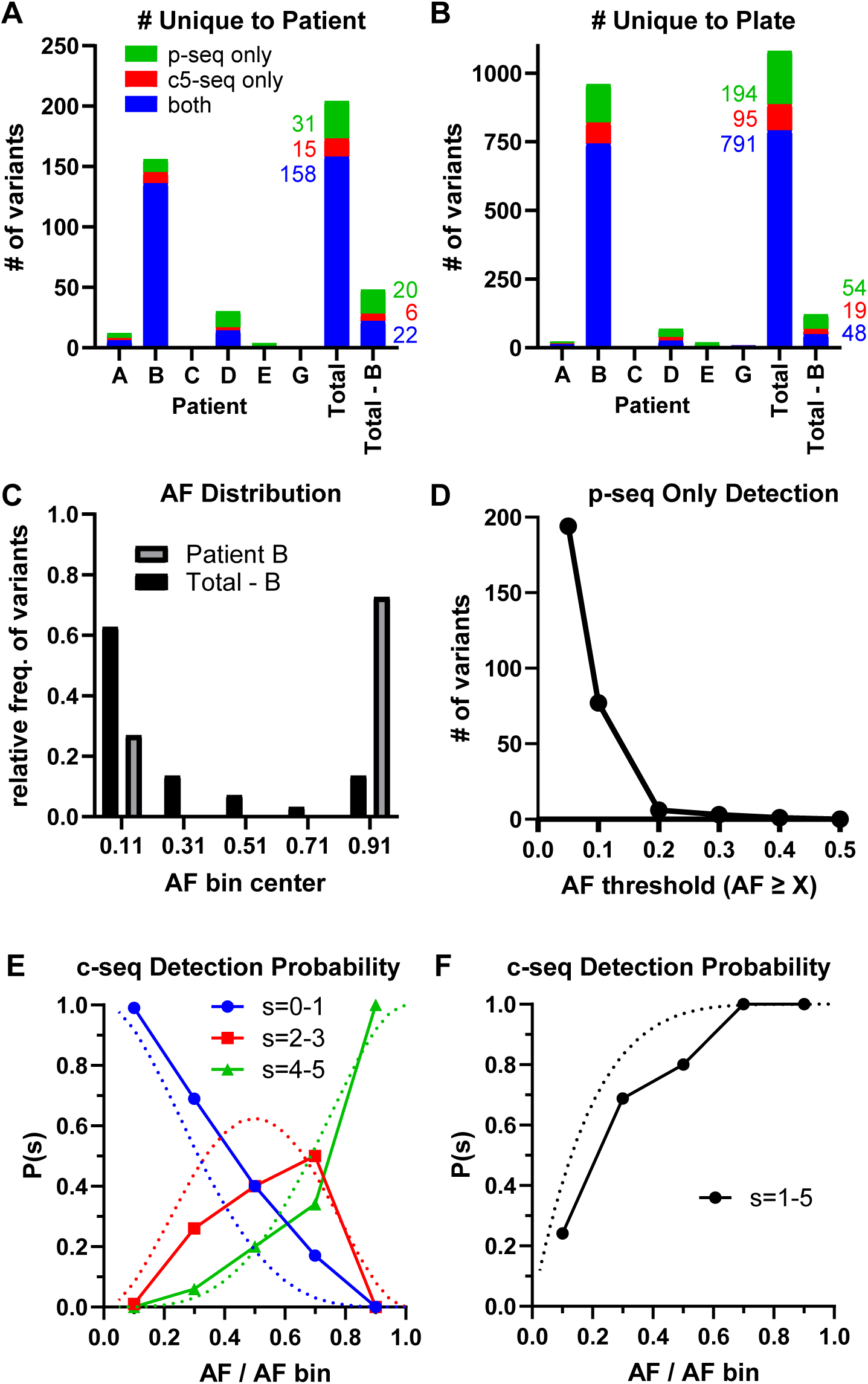
Sensitivity of variant allele detection. **A.** Stacked bar plot of the number of variants for each detection modality when counting those unique to each patient. **B.** Stacked bar plot of the number of variants when counting those unique to each plate (allows variants to be counted multiple times if they were on more than one plate). **C.** Allele frequency distribution of p-seq data from Patient B or all patients combined (excluding Patient B). **D.** Number of variants detected only by p-seq (and not c5-seq) as a function of allele frequency (AF) inclusion threshold. **E.** Probability of c5-seq variant detection as a function of p-seq AF. The observed detection probabilities for c5-seq were calculated by grouping variants from each blood culture plate into p-seq AF bins, then calculating the proportion of the five colonies in the c5-seq dataset that detected each allele in that bin. Data symbols indicate the observed probability for each AF bin (plotted at the bin centers) and are connected by solid lines. The AF bin ranges were 0.05≤x≤0.2, 0.2<x≤0.4, 0.4<x≤0.6, 0.6<x≤0.8, and 0.8<x≤1.0. Dotted lines indicate the theoretical probability plotted as a continuous function based on the binomial model. For example, P(s=0-1) is the probability that 0 or 1 colony (out of the 5 colonies sampled from the plate) detected the variant. **F.** P(s=1-5) is the probability that any of the five colonies sampled detected the variant allele (i.e. the overall sensitivity of c5-seq).

However, we noted that most variants from this six-patient dataset were derived from Patient B alone. This patient (discussed in detail below) was infected with two ‘clades’ of clonal variants that differed by as many as 99 SNPs (**Fig. S1**). During this infection episode, clones from one clade completely displaced the other clade. As a result, all clade-distinguishing variants rose to AF=1. This large number of high frequency variants from Patient B skewed the AF distribution in the overall dataset (**Fig. 3C**) and our analysis showed that Patient B was an outlier in this respect. To obtain a more representative comparison of p-seq and c5-seq, we removed Patient B and repeated the above analysis (**Fig. 3A**, see “Total – B”). Now, out of 48 total variants, p-seq detected 44 (92%) and c5-seq detected only 28 (58%), yielding p-seq 1.6-fold (44/22) more variants than c5-seq. We also analyzed these data with Patient B removed on a per plate basis (**Fig. 3B**, see “Total – B”) and this resulted in a similar distribution. Out of 121 total variants, p-seq detected 102 (84%) and c5-seq detected 67 (55%), yielding p-seq 1.5-fold more variants than c5-seq. We conclude that sensitivity can differ widely between patients based on the specific AF distribution of the infection and that p-seq has a variant detection rate that is ∼1.5-fold greater than that of c5-seq.

We next sought to determine the plate AF range where p-seq offered more sensitivity than c5-seq and further characterize the false-positive rate for p-seq. First, we plotted the number of variants in the entire dataset detected only by p-seq as a function of p-seq AF threshold (**Fig. 3D**). We found that p-seq only offered significant additional variant detection sensitivity over that of c5-seq for AF <0.2. We reasoned that false-positive detections should therefore also be restricted to AF <0.2. Therefore, we counted the number of p-seq samples that resulted in zero variants detected only by p-seq with AF <0.2, since a zero result in this range also equates to zero false-positive detections. Of the 107 samples we analyzed by p-seq, notably, 77 (72%) samples were zeros. This shows our false-positive rate was very low across the dataset. Even if we conservatively assume that the other 30 (28%) plates each had one false-positive, this places an upper bound of ∼0.3 (30/107) false detections per sample, which is consistent with the estimated rate of 0.2 per sample from our control dataset. In the six-person dataset discussed above, after analyzing 75 blood culture plates with both p-seq and c-seq, 194 variants were discovered by p-seq only. We estimate that <23 (75 samples x 0.3 per sample) of these are false positives.

We next aimed to characterize c5-seq detection sensitivity. At higher variant AF, a higher proportion of the five sampled colonies should contain the variant. To examine the performance of c5-seq variant allele detection, we plotted the probability (P) that 0 to 1 (s=0-1), 2 to 3 (s=2-3), or 4 to 5 (s=4-5) colonies would be positive for the variant allele as a function of the p-seq variant AF. We found the probability of detection by c5-seq generally agreed with the theoretical probabilities modeled by the binomial distribution (**Fig. 3E**). This analysis confirms the expectation that the proportion of colonies positive for a given variant allele approximates the frequency on the plate. This is important because it rules out the possibility that major systematic errors are introduced when choosing five isolated colonies for sequencing. Finally, we assessed the overall sensitivity of c5-seq by plotting the probability that any of the 5 colonies contained the variant allele (s=1-5) (**Fig. 3F**). This result agrees well with the binomial model and shows that the overall detection rate of c5-seq approached 100% for AF>0.6 and was 24% for AF<0.2.

### Within-host genetic variation

To characterize the polyclonal variation within each patient, we used the c5-seq dataset to count the number of unique genotypes on each of the 75 blood culture plates (**Fig. 4A**). 48% of the plates were polyclonal (i.e. >1 genotype) and all patients had at least one polyclonal plate. The plates from Patients C, G and E were relatively less diverse, with 1-2 genotypes/plate and most with only one genotype. In contrast, the plates from Patients A, B, and D contained 1-4 genotypes/plate, with most plates having >1 genotype. For these latter patients, the greater number of polyclonal plates was not merely due to repeated identification of the same genotypes. Rather, these three patients each had more diverse infections overall, each with ≥10 total unique genotypes identified (**Fig. 4B**).

**Figure 4.**
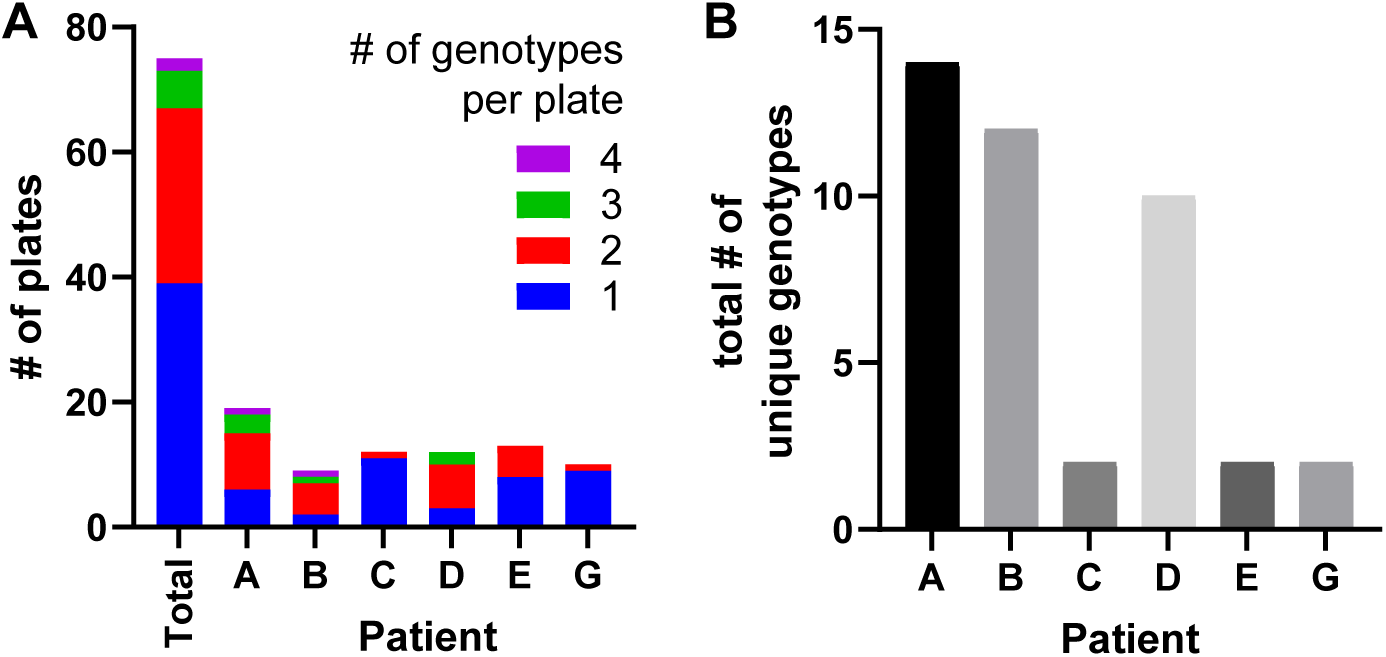
Polyclonal variation. **A.** Stacked bar plot of the number of plates analyzed for each patient, or all the patients (Total). The number of genotypes per plate is indicated by different colors. **B.** Bar plot of the total number of unique genotypes identified for each patient.

Blood was sampled longitudinally over several days for each patient. To understand how this type of repeat sampling impacted the detection of new variant alleles, we constructed variant accumulation curves for each patient by plotting the cumulative number of unique variant alleles detected as a function of sampling day. Plots of the c5-seq data (**Fig. 5A**) and p-seq data (**Fig. 5B**) were similar and revealed two distinct patterns. For seven of the patients (Patients C and E-J), longitudinal sampling added ≤3 new alleles and the number of new alleles detected plateaued with time, suggesting sampling approached saturation. In contrast, for the three patients we identified above with highly diverse infecting populations (Patients A, B, and D), new variants continued to be detected throughout the entire sampling period. This pattern shows that there is greater variant richness for these three patients and ongoing evolution of genotypes.

**Figure 5.**
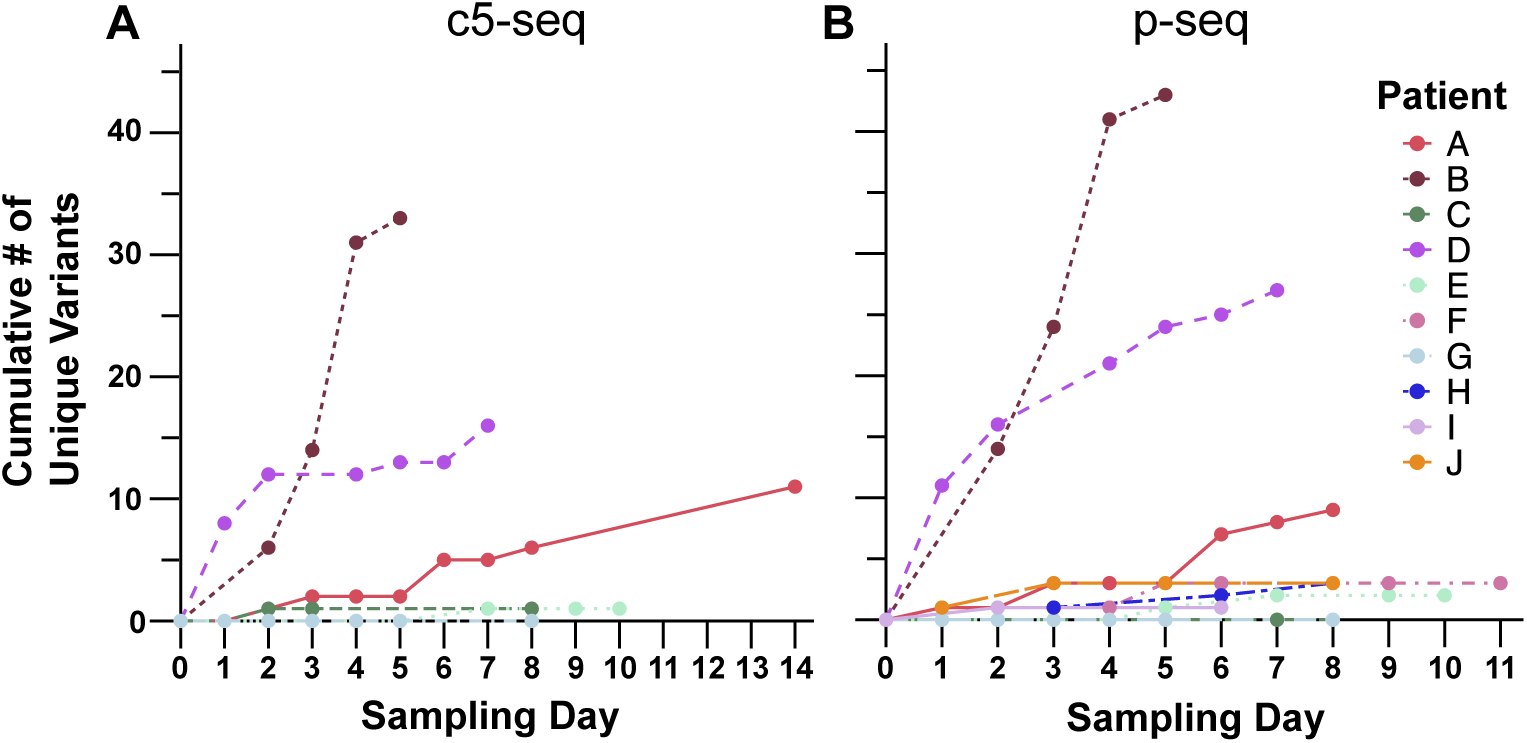
Variant accumulation curves. The cumulative number of unique allelic variants identified for each patient by (A) c5-seq or (B) p-seq is plotted as a function of sampling day. For ease of visualization, each curve is normalized to start at zero unique variants. This was achieved by subtracting the number of unique variants sampled on day 0 from all values of the curve.

### Evolutionary dynamics

To visualize evolutionary dynamics in the three patients with more diverse infections, we used the c5-seq data to construct Muller plots of genotype frequency as a function of time (**Fig. 6**). For most of the sampling timepoints we applied c5-seq to two blood culture plates, so ten genotypes per timepoint were usually used to calculate genotype frequencies (**File S1**). Given the importance of antibiotic pressure in facilitating evolutionary dynamics, we overlayed the time periods of anti-MRSA antibiotic exposures onto these plots. Major shifts in genotype frequency were associated with a change in specific antibiotic exposures, as detailed for each patient below:

**Figure 6.**
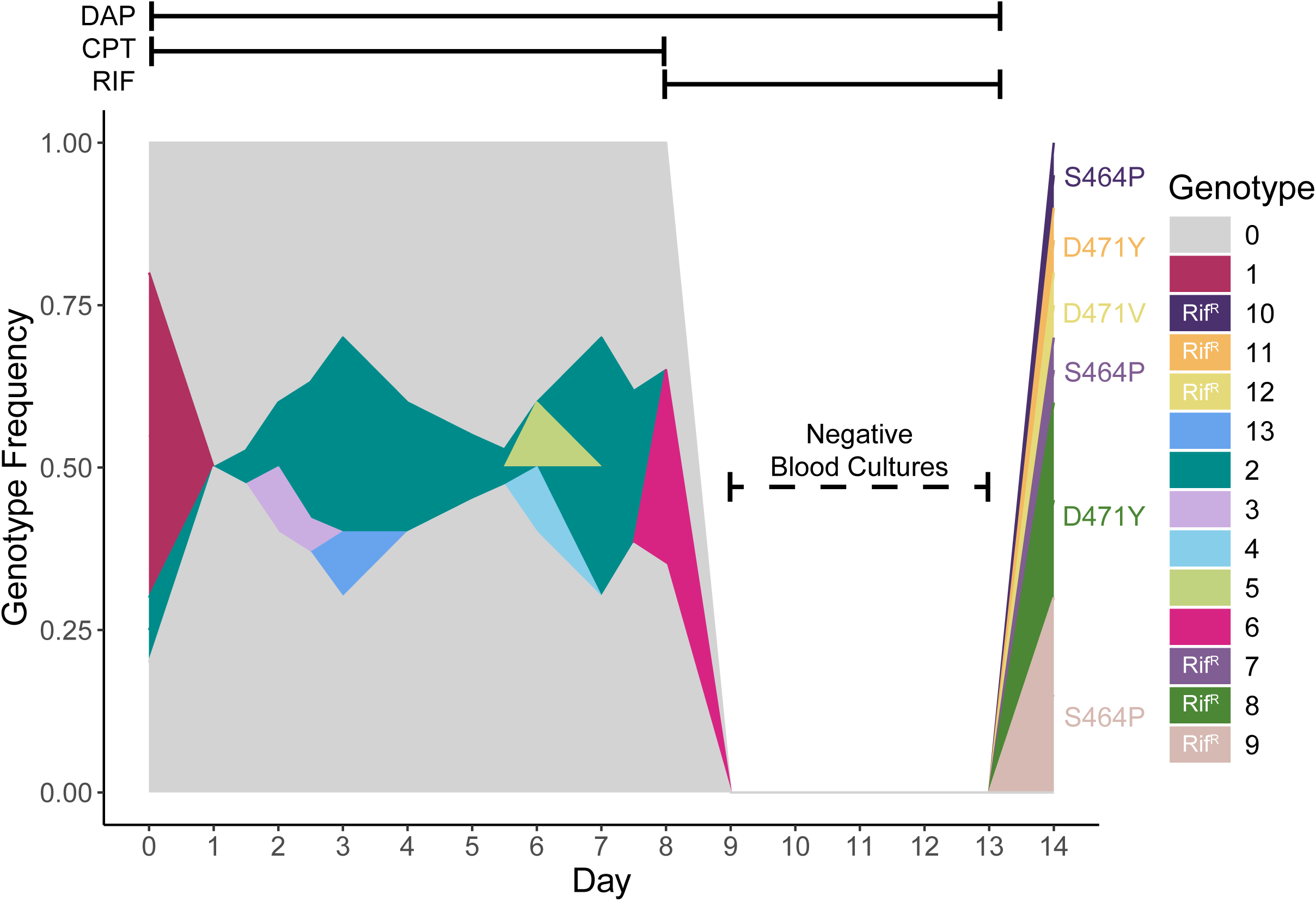
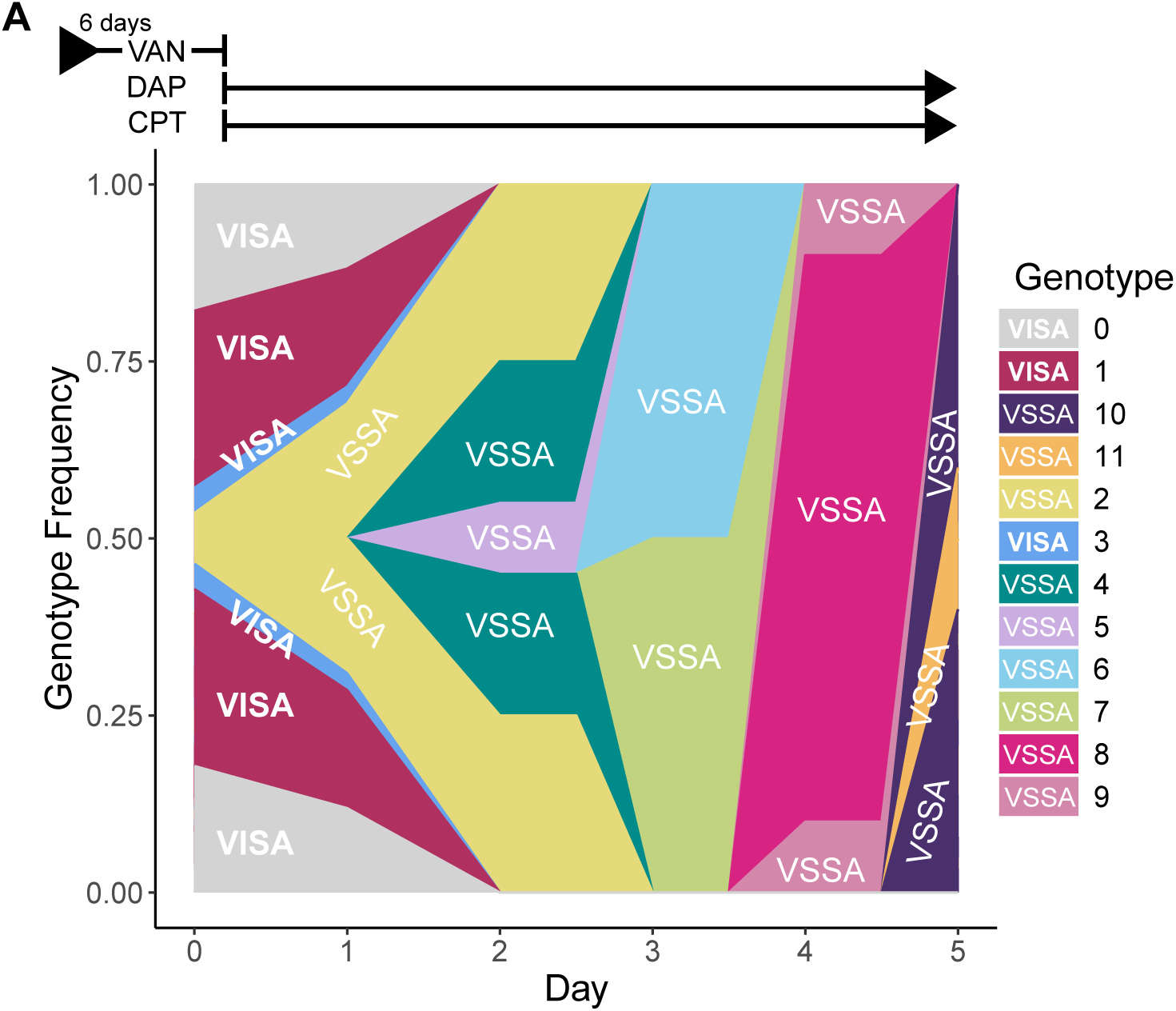
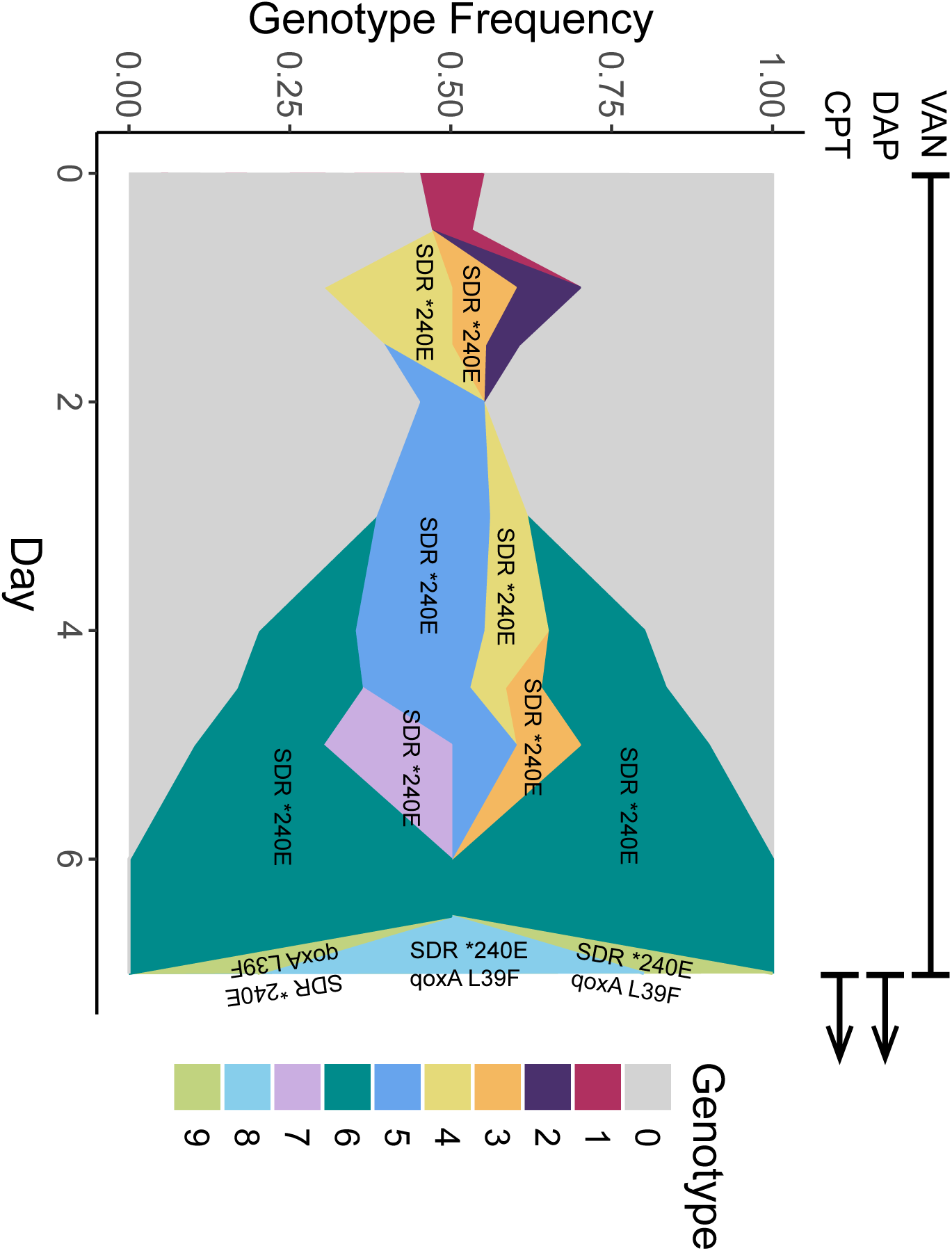
Muller plots. **A.** Patient A. Genotypes that evolved resistance to rifampin (Rif^R^) are annotated in the legend with the specific RpoB amino acid substitutions noted next to each genotype plot. **B.** Patient B. Genotypes in the vancomycin-intermediate *S. aureus* (VISA) and vancomycin-susceptible *S. aureus* (VSSA) clades are annotated in the legend and labeled in the plot. **C.** Patient D. Genotypes with mutations in the short-chain dehydrogenase/reductase (SDR) family protein of unknown function (SDR *240E) and *qoxA* (*qoxA* L39F) are labeled in the plot. Time periods of anti-MRSA antibiotic exposures are indicated above each plot (CPT=ceftaroline, DAP=daptomycin, RIF=rifampin, VAN=vancomycin). See supplementary data for the complete list of mutations present in each genotype (**File S1**).

#### Patient A

Patient A was diagnosed with prosthetic valve endocarditis and was treated with antibiotics without valve surgery. Despite 1 week of combination therapy with daptomycin and ceftaroline, blood cultures remained positive. On day 8, ceftaroline was stopped due to possible toxicity, daptomycin was continued, and rifampin was added as salvage therapy. This change in antibiotics was associated with bacterial clearance from the blood, as blood cultures obtained on days 9, 10, 11, and 12 showed no growth. Unfortunately, the patient died on day 13. The following day, one set of blood cultures was obtained at autopsy, and both bottles were positive for MRSA. Six new genotypes were identified in this culture (**Fig. 6A**). Notably, all ten clones sampled on day 14 encoded one of three different *rpoB* mutations known to confer resistance to rifampin (S464P, D471Y, D471V) (10, 11) and exhibited phenotypic resistance to rifampin by disc diffusion assay (**Table S2**). *rpoB* encodes the β subunit of RNA polymerase, which is the target of rifamycin antibiotics. Rifampin exposure during concomitant *S. aureus* bacteremia is associated with the rapid acquisition of resistance even when used as adjunctive therapy (12). This analysis shows that resistance evolved independently at least three times within six days of the initial drug exposure in this patient. These data support the notion that when adjunctive rifampin is indicated for an infection due to staphylococci, clinicians should consider delaying its initiation until the burden of infection is lowered to prevent the emergence of resistance (13).

#### Patient B

Patient B was diagnosed with prosthetic valve endocarditis complicated by paravalvular abscess four months prior to arrival at our institution and had been treated with antibiotics without valve surgery. Treatment constituted a 3-month course of combination therapy with vancomycin and rifampin that included gentamicin for the first 2 weeks, then antibiotics were discontinued. Three weeks later, the patient was diagnosed with recurrent bacteremia at another hospital and received six days of vancomycin prior to transfer to our institution. Only the blood cultures that were collected after arrival at our institution were available for this analysis. Interestingly, we found this patient was infected with clonal variants that differed by as many as 99 SNPs, and phylogenetic analysis showed that variants grouped into two distinct clades (**Fig. S1**). MIC tests of representative isolates for each of the 12 different genotypes revealed one clade (genotypes 0, 1, and 3) was comprised of vancomycin-intermediate *S. aureus* (VISA) and the other clade (genotypes 2 and 4-11) was comprised of vancomycin-susceptible *S. aureus* (VSSA) (**Table S2**).

Shortly after the initial day 0 blood cultures were collected, the patient was switched from vancomycin to combination therapy with daptomycin and ceftaroline, which coincided with a major shift in genotype frequency (**Fig. 6B**). The day 0 samples contained 4 different genotypes; VISA clones (genotypes 0, 1, and 3) constituted 93% of the population, whereas the VSSA clone (genotype-2) constituted only 7%. However, by day 2, and coinciding with cessation of vancomycin, VSSA clones increased in frequency to became 100% of the population, thereafter, completely displacing the VISA clones. These data suggest that removal of vancomycin from the antibiotic regimen played a major role in permitting the VSSA clones to sweep the population.

The relatively large genetic distance (99 SNPs) separating the VISA and VSSA clades in this patient is notable, as most clones isolates from the same episode of *S. aureus* bacteremia differ by <10 SNPs (3, 14). Although we found that both clades belonged to the same multi-locus sequence type, ST72, we considered the possibility that two closely related, yet distinct, strains independently caused invasive infections. While this is possible, mixed strain invasive infections are exceedingly rare for *S. aureus* (4). We also considered acquisition of a hypermutator phenotype. However, we did not detect mutations in genes associated with hypermutation in *S. aureus* (*mutL* and *mutS*) and did not observe colonies within the zone of inhibition in our rifampin disc diffusion assay that would be characteristic of hypermutation. Given the clinical history of 4 months of persistent infection, an alternative hypothesis is that the VISA and VSSA clades diverged from a common ancestor after prosthetic valve endocarditis was established. In support of this idea, both clades share the identical and relatively rare N474K rifampin resistance mutation in *rpoB* (15), which suggests genetic divergence occurred after the patient’s exposure to rifampin. We speculate the clades evolved in distinct niches with differential exposure to vancomycin (e.g. inside versus outside the valve abscess).

#### Patient D

Patient D was a severely immunosuppressed patient due to a recent solid organ transplant complicated by acute graft rejection and was diagnosed with upper extremity cellulitis and septic pulmonary emboli. Transesophageal echocardiography was negative for endocarditis, suggesting the valvular vegetation was either small, had completely embolized, or that the source of septic pulmonary emboli was an upper extremity septic thrombophlebitis. Despite 7 days of vancomycin, blood cultures remained positive and so the patient was switched to a combination of daptomycin and ceftaroline on day 7. We observed that a new genotype (genotype-6) emerged by day 4 and steadily rose to overtake the population on day 6 (**Fig. 6C**). Genotype-6 contains a mutation in a gene encoding a protein of unknown function that is conserved in *S. aureus* and annotated as a short-chain dehydrogenase/reductase (SDR) family NAD(P)-dependent oxidoreductase (NCBI RefSeq: WP_000678232.1). Relative to the gene sequence of the earlier isolates, the mutation causes a base pair substitution within a stop codon at codon position 240 conferring a change to glutamate (*240E). Considering the location of the next in-frame stop codon, this results in an extension of the protein sequence by 12 residues. BLAST searches (16) of these two protein sequences against the NCBI ClusteredNR database (version: 2026/03/14) revealed heterogeneity in the gene length of sequenced *S. aureus* strains, including multiple examples identical to the two variants in this patient. The significance of this variation is unclear but argues against a major functional impact of the *240E variant. We also noted that the change in antibiotics on day 7 was associated with two new genotypes that swept the population and contained additional mutations on the background of genotype-6. Both new genotypes evolved a mutation in *qoxA* conferring an L39F amino acid substitution that maps to a transmembrane domain. The *qoxA* mutation may be significant, as the *qoxABCD* operon encodes one of the two main terminal oxidases needed for respiration in *S. aureus* (17) and loss of respiratory capacity is associated with persistent infection and antibiotic tolerance. Protein-altering mutations in *qoxA*, *qoxB*, and *qoxD* have also been identified in other clinical *S. aureus* isolates (3).

### Evidence of parallel evolution

Independent mutation of the same genes or pathways can be an indication of parallel evolution, suggesting the same adaptation evolved in different infections in response to a similar environmental stress. Three alleles (*rpoB*, *pdxK*, and *glcB*) in our dataset evolved multiple independent protein-coding mutations, and so we examined the literature for further evidence to support parallel evolution in each case. Mutations in *rpoB* are relatively common and represent a known genetic signature of within-host evolution for *S. aureus* (3). In our dataset, in addition to Patient A (discussed above), Patient B evolved a mutation in *rpoB* causing an R155C amino acid substitution that rose to fixation. This mutation occurred on the background of an N474K mutation already present in all isolates recovered from Patient B. The N474K mutation falls within the rifampin-resistance determining region (RRDR) of *rpoB* and is known to cause resistance (15), and likely evolved in response to the patient’s prior rifampin exposure. In contrast, the R155C mutation lies outside the RRDR and we found it did not further alter rifampin susceptibility (**Table S2**). The significance of this mutation is unclear, although one hypothesis is that it is a compensatory mutation. For example, for *Mycobacterium tuberculosis*, *rpoB* mutations outside the RRDR have been described that alleviate fitness defects caused by rifampin-resistance mutations (18). *rpoB* mutations are also known to alter global transcription patterns that affect other bacterial phenotypes that may be relevant *in vivo* (19, 20). Next, *pdxK* encodes a pyridoxal kinase that is required in the vitamin B6 salvage pathway of *S. aureus* (21). Patient B evolved a mutation conferring an A122T amino acid substitution that rose from <10% to 100% of the population, and Patient H evolved a mutation conferring an A180T amino acid substitution that rose from 0% to 84% of the population. The possible functional impacts of these substitutions on pyridoxal kinase activity are not clear, however, we noted that A180T maps to a conserved ‘GG switch’ region near the ATP binding pocket (21), and thus could possibly alter enzyme function. Protein-altering mutations in *pdxK* have also been identified in other clinical *S. aureus* isolates (3). Finally, *glcB* encodes a glucose-specific EIICBA component of a phosphotransferase system for sugar import. Patients F and H both evolved the same mutation conferring an M509L substitution, which resides within a hydrophobic core of the protein. Although the probability of detecting independent identical base pair substitutions by chance alone is very low, we also noted in this case that both variants were low frequency (AF = 0.05 - 0.06), were detected from a single sampling plate, and that the amino acid substitution conferred is physicochemically conservative. Ultimately, additional experiments would be needed to test for a functional impact and a possible selective advantage for all three of these variant alleles to demonstrate parallel evolution.

We also reviewed our dataset for singleton mutations in genes and pathways that have previously been identified as hot spots for parallel evolution in *S. aureus*. Giulieri, *et al*. published a detailed gene enrichment analysis using 2,590 *S. aureus* genomes derived from 396 independent episodes of infection (3). We found mutations in *saeS* and *vraS*, which are two of the genes identified in their study. Additionally, we noted the absence in our dataset of commonly mutated genes. These included the *agr* operon (virulence), *mprF* (daptomycin resistance), *stp1* (vancomycin resistance), and tricarboxylic acid (TCA) cycle genes (antibiotic tolerance). In a study we published previously that analyzed 206 MRSA isolates derived from 20 independent episodes of infection, we also identified mutations in *mprF* and TCA cycle genes (14). In that study, we sequenced only one colony per each of the 206 blood culture plates. Here, deeper sampling of 107 plates with p-seq did not significantly improve the identification of the most common genetic signatures of parallel evolution.

## DISCUSSION

The optimal approach to using bacterial WGS to characterize genetic diversity in clinical blood culture specimens is unclear. To understand this further, we analyzed the blood culture plates from cases of persistent MRSA bacteremia with both p-seq and c-seq and our analysis revealed the tradeoffs of these two different methods. As expected, the primary advantage of p-seq was that it is more sensitive than c5-seq for detecting low frequency variant alleles. It is also noteworthy that p-seq is the more economical method because it requires only a single DNA library preparation for WGS per blood culture plate, whereas c5-seq, for example, incurs the costs associated with five separate library preps. However, we found p-seq also has two main disadvantages that need to be weighed against its greater sensitivity.

First, variant detection with next generation sequencing data is fraught with low frequency false-positive variant calls due to errors that arise from library preparation, PCR, sequencing, and read alignment (22) and this is a much more prominent issue with p-seq. With c-seq, this problem is mostly eliminated because the goal is to find the consensus genome sequence of the pure colony, so these low frequency false-positive variants are easily excluded with a relatively high (more stringent) consensus AF cutoff value (e.g. AF≥0.8). This includes filtering *de novo* mutations that arise *ex vivo* during colony formation, which are also expected to be low frequency and are false-positive in this context. In contrast, p-seq aims to characterize a mixture of different genome sequences, so detecting lower frequency true-positive variants is the objective. However, distinguishing between true-positive and false-positive variants is more challenging and mandates read alignment to a high-quality reference genome, manual data inspection, and custom data filtering, which are not standardized and often vary across variant callers and research groups. In our study, we chose to validate our filtering approach using a control dataset and estimating the false-detection rate, however, this adds time and cost to the analysis. We found that p-seq had a 1.5-fold higher variant detection rate than c5-seq. Although our false-detection rate was quite low (<0.3 per sample), a subset of variants are still likely false positives. Note that we chose to filter out variants with AF <0.05 in our study, but to more stringently mitigate false-positive variant calls, investigators may instead choose to raise the AF filtering cutoff (e.g. AF <0.2). Indeed, assuming read quality metrics are incorporated as a filtering strategy (we used bias p-value scores reported by *breseq*) (**Fig. 2B**), this would essentially eliminate false-positives. However, we found that p-seq only offered additional variant detection beyond that of c5-seq for AF <0.2 (**Fig. 3B**). Thus, while raising the AF cutoff is a valid filtering strategy for p-seq, in our dataset, this no longer provided any variant detection sensitivity advantage over c5-seq, thereby negating the primary technical advantage of the method.

Second, when bacteria are pooled prior to sequencing, genetic linkage information is lost. c-seq returns a specific genotype for each colony isolate, whereas p-seq returns only a list of allele frequencies from a pooled mixture. This makes tracking the evolution of genotypes a challenge with p-seq. While it is possible to reconstruct genotypes from population-level sequencing data that is collected longitudinally (23), this requires relatively deep sampling of the evolving bacterial population at high temporal resolution and is generally more feasible for analyzing laboratory-based evolution experiments than for clinical samples. For blood cultures, there is expected to be a small sampling bottle neck size (discussed below) and our data indicated rapid shifts in genotype frequencies between time points. These features would make genotype reconstruction from p-seq less accurate and informed our decision to construct Muller plots using the c-seq dataset. An additional disadvantage of p-seq is that subsequent genotype-phenotype studies are hindered because pure clones with known genotypes are not readily available from the cryopreserved stock of a pooled mixture. For example, we were only able correlate antibiotic resistance phenotypes to specific genotypes for patients because we had c-seq data for each individually cryopreserved clone. Thus, depending on the application, utilizing c-seq, p-seq, or a combination of the two may be optimal.

We suspect our findings can be generalized to blood cultures of other common bacterial pathogens but may be less relevant for other specimen types. The most critical parameter to consider is the expected sample bottleneck size, as the utility of p-seq diminishes with smaller sample populations. For example, in the extreme case of a 1 CFU sample bottleneck, the theoretical sensitivity benefit of p-seq over c-seq is rendered completely moot. Most clinical specimens are directly streaked onto culture plates and so the number of CFUs yielded reflects the size of the bottleneck. For example, p-seq has been used to analyze MRSA nasal surveillance cultures where culture plates were directly inoculated with nasal swabs and yielded 8 to 200 CFU for each pool (2). The authors of that study suggested investigators could define a minimum pool size in which to deploy p-seq. For specimens that are directly streaked onto culture plates, one can simply count the number of CFUs present on the plate to decide if it warrants deep sequencing by p-seq. However, this approach is not possible for blood cultures since the patient’s blood specimen is first used to inoculate liquid broth and bacteria are grown in culture prior to plating. It is estimated that 50% of bacteremia cases have a bacterial blood concentration in the lower range of 0.01 - 1 CFU/mL, and may only rarely exceed 100 CFU/mL at the high end of the range (7). To detect these low pathogen concentrations, blood culture bottles containing nutrient rich broth are inoculated with ∼10 mL of patient blood (1 – 1000 CFU) and may be incubated for up to 5 or more days to facilitate bacterial growth (**Fig. 1B**). Two to three sets of blood cultures bottles (4 to 6 bottles, or ∼40-60 mL of blood total) are required to increase the sensitivity of detecting bacteremia to >90% (24). Bottles are deemed “positive” if bacterial growth occurs, at which point the bottle contains 10^8^ - 10^9^ CFU/mL. Then, a liquid drop (>10^5^ CFU) of the bottle culture is used to inoculate solid media (i.e. blood culture plate) via steak-plating to obtain isolated colonies. Therefore, although the blood culture plate is inoculated with >10^5^ CFUs, there is an initial sampling bottleneck that is often only 1 – 10 CFU. This is likely a major reason why p-seq only offered a modest increase in sensitivity over c5-seq in our study. More importantly, however, we found that analyzing all the available blood cultures from each episode of bacteremia was critical for identifying and characterizing genetically diverse infections. Simply put, more blood is better. Our data show that even when using a p-seq strategy with optimized sensitivity, the major limitation for characterizing diversity in blood cultures becomes the number of blood cultures available for analysis. It should be noted that the sampling bottleneck also places a limitation on accurately characterizing evolutionary dynamics too. Small sampling bottlenecks can artificially inflate variant frequency by the random process of genetic drift via the founder effect. In our dataset, we were able to mitigate this issue when constructing Muller plots because we generally had multiple positive blood cultures at each time point and plotted the average genotype frequency.

Our analysis revealed detailed examples of rapid evolutionary dynamics as well as signals of evolution by selection. Curiously, we also noted that the variant AF distribution on blood culture plates was bimodal and heavily skewed toward AF ≥0.9 and AF <0.3 (**Fig. 3C**). Such U-shaped or J-shaped skews in AF distributions could be caused *ex vivo* by artificially imposed genetic drift due to the sampling bottleneck mentioned above, however, there are several factors *in vivo* that can also lead to rapid sweeps of specific alleles to high frequency. For example, in murine models of *S. aureus* bacteremia, individual bacteria seed tissues and form microabscesses by clonal expansion, of which only some abscess populations survive antibiotic killing in a stochastic manner (25, 26), a mechanism that facilitates evolution by genetic drift. Also, clones carrying strongly beneficial mutations may rise to high frequency quickly. For example, Patient A evolved rifampin resistance mutations independently at least three times after exposure to the drug. Additionally, neutral variant alleles that are linked to a beneficial mutation on the same genome can also rise to high frequency, a phenomenon referred to as genetic hitchhiking. For example, Patient B was infected with two distinct clades that differed by ∼100 variants. The clones of the VSSA-clade swept an infecting population dominated by clones of the VISA-clade after vancomycin was stopped. This resulted in ∼100 variants rapidly rising to fixation in the population (AF=1) at later times, leading to a marked ‘J-shaped’ skew in the AF distribution (**Fig. 3C**). We also identified possible signatures of parallel evolution by identifying genes that were repeatedly mutated in this and other studies (e.g. *rpoB*, *pdxK*, *qoxA*, *saeS*, and *vraS*). Importantly, however, we found that deep sampling with p-seq did not identify many of the most common mutated genes in *S. aureus* reported in the literature. Thus, p-seq does not remove the requirement to analyze large numbers of bacteremia episodes to identify common genetic signatures of parallel evolution.

Our data suggests a more cost-efficient method to track evolution for cases of persistent bacteremia. We found that sampling blood culture plates from several time points is critical, but only for the subset of patients that have especially diverse infections. Notably, analyzing only the first and last positive blood cultures in some of the cases we analyzed would have missed a significant amount of diversity, as alleles can rise above and then fall below the detection limit in between sampling timepoints. On the other hand, most of the infection episodes in our dataset were not genetically diverse and had ≤3 total variant alleles. In these cases, applying c5-seq to all the plates did not yield much additional information beyond what could have been generated from the analysis of a single sample. This suggests an approach that leverages both c5-seq and p-seq. In this hybrid approach, the investigator still samples and archives individual colonies and entire plate populations, but then first uses the more economical p-seq to screen for diversity on all the plate samples. Then, once the diverse infections (or plates) are identified, c-seq can be applied to only the samples with high genetic diversity to obtain specific genotypes. For example, if we had applied this procedure to the six patients we analyzed with both c5-seq and p-seq it would have saved us the cost of performing c5-seq on 47% of our total samples (175 of 375 total colonies) and, notably, no mutations would have been missed.

Three of the ten patients in our study (patients A, B, and D) had especially genetically diverse infections. Rarely, *S. aureus* can evolve a hypermutator phenotype in the setting of chronic infection (27). Hypermutator strains have mutation rates ∼100 times greater than baseline and will form colonies within the zone of inhibition on rifampin disc diffusion assays. However, we did not observe colonies within the inhibition zone for these isolates in rifampin disk diffusion assays that would be indicative of hypermutation, and we also did not detect mutations in genes known to cause the hypermutator phenotype in *S. aureus* (*mutL*, *mutS*). Instead, our data point to patient-level factors expected to be associated with polyclonal bacteremia. Patients A and B were the only two patients with prosthetic valve endocarditis, and Patient D had an endovascular infection in the setting of severe immunosuppression. These types of infections have a persistent high bacterial burden that is in direct continuity with the bloodstream. These features would be expected to facilitate both the evolution and subsequent detection of variants. Future studies of within-host bacterial evolution may consider specifically targeting this patient population for study and use the hybrid sequencing approach suggested above for a cost-efficient discovery and analysis of variants.

In conclusion, our study shows the strengths and limitations of colony-only versus pooled-sequencing. Although pooled sequencing is more cost-effective and enables the detection of lower frequency variants, the need to mitigate false-positive variant calls and the lack of genotypic information were major disadvantages. In addition to comparing these methods, we were able to characterize evolutionary dynamics in response to antibiotic exposure and find evidence suggesting parallel evolution across patients. The prevalence and generalizability of these latter findings are limited by our study design since we only included ten patients with persistent MRSA bacteremia at a single healthcare center, which likely does not accurately represent the diversity or variant dynamics across a larger patient population. Although this study focused on MRSA bacteremia, the findings are likely to be directly pertinent to other bacteremia pathogens. While we caution extrapolation of our findings to other culture specimen types, the evaluation of methods described are likely to be broadly useful for these investigations.

## Materials and methods

### Clinical cases

Patients admitted to the University of Pittsburgh Medical Center with MRSA-PB were identified by consecutive dates of positive blood cultures over a period spanning from January 2020 to September 2021. Patients were selected based on prolonged bacteremia, which ranged in duration from 6 to 15 days. Patient characteristics are given in **Table S1**.

### Blood culture plate sampling, isolate storage, and DNA isolation

The agar plates (blood or chocolate agar) that were used to streak out positive blood cultures from each patient were collected after colony formation and sampled. For c5-seq, each single colony isolate was cultured overnight in tryptic soy broth (TSB) and archived in 15% glycerol at -80°C. Then, for each archived colony, genomic DNA (gDNA) was extracted from an overnight culture inoculated from the archived glycerol stock. For p-seq, each plate was washed with 5 mL of TSB until all visible bacterial growth was in a pooled suspension, which was directly archived in 15% glycerol at -80°C without subculture. Then, for each plate pool, gDNA was extracted directly from the archived pool without subculture. gDNA was extracted using the DNeasy Blood and Tissue Kit (Qiagen) using the manufacturer protocol, except that lysostaphin (Sigma) was added to facilitate lysis of *S. aureus*. DNA was quantified using a NanoDrop instrument (Thermo Fisher Scientific). Samples submitted for DNA sequencing had concentrations ≥30 ng/μL. For the experiment that analyzed variant detection using a mixture of gDNA from two different *S. aureus* clones, gDNA was quantified using a Qubit Fluorometer (Thermo Fisher Scientific), then mixed at known ratios (50:50, 80:20, 90:10, 95:5, and 99:1) keeping the total DNA concentration fixed at 50 ng/μL.

### WGS

Illumina DNA sequencing libraries were constructed using Nextera DNA kits (Illumina) with each isolate assigned a unique barcode. Sequencing was carried out on the NextSeq 2000 platform (Illumina). gDNA isolated from individual clones (c-seq) was sequenced to an average depth of 154X ± 43X (mean ± SD). gDNA from plate populations (p-seq) was sequenced to an average depth of 710X ± 196X (mean ± SD). For the experiment that analyzed variant detection using a mixture of gDNA from two different *S. aureus* clones, the mixtures were sequenced to an average depth of 1073X ± 44X (mean ± SD). Sequencing data derived from one individual colony was used to construct a custom reference genome for each patient. To obtain a high-quality reference, this DNA was also sequenced using the MinION platform (Oxford Nanopore Technologies), then genomes were assembled *de novo* by combining the Illumina and MinION data (28) via Unicycler v0.5.1 (29). Prokka v1.14.5 (30) was used for annotation. MinION libraries were prepared using a rapid barcoding kit (SQK-RBK004) and sequenced on R9.4.1 flow cells. Albacore v2.3.3 or Guppy v2.3.1 were used for the base-calling on raw reads (Oxford Nanopore Technologies).

### Variant identification and filtering

Illumina reads were analyzed with *kraken 2* (9) to look for sample contamination from other bacterial species that could interfere with variant calling. We detected *Escherichia coli* in two p-seq samples from Patient J and we detected *Micrococcus luteus* in one c-seq sample from Patient B and so did not include these samples in our analysis. DNA sequence variants were identified with *breseq* v0.39.0 (8) using the custom genome derived from the same infection as the reference. For c-seq, *breseq* was run in ‘consensus mode’ using default parameters (e.g. consensus frequency cutoff = 0.8). For p-seq, *breseq* was run in ‘polymorphism mode’ using an AF threshold of 0.05, then variant calls were further filtered as described above (see Results). Variants that passed this filter had a minimum depth of coverage of 21 and at least 11 reads supporting the variant-call. For the experiment that analyzed mixtures of gDNA of two different clones at known ratios, the AF threshold in *breseq* was adjusted down to 0.01 for the 95:5 ratio and adjusted down to 0.001 for the 99:1 ratio.

### Phylogenetic analysis of Patient B isolates

Illumina reads from c-seq data were aligned to the reference genome using *snippy* v4.6.0 (31). The alignment was stripped of gaps and identical sites in Geneious v2023.1.1. The phylogenetic tree was constructed in Geneious using the UPGMA model with 1000 bootstraps and visualized in iTOL v7(32).

### Antibiotic susceptibility testing

MIC and disk diffusion assays were performed and interpreted per Clinical Laboratory Standards Institute guidelines (33). MICs were determined by broth microdilution in 96-well plates. Reported MIC and zone of inhibition diameter values are the average of at least two independent replicates.

### Statistics and data visualization

Data plots and histograms were generated using R or Prism (GraphPad Software). To estimate the error of p-seq allele frequencies, the mean absolute difference was calculated using the average of all twenty-five values for |AF_obs_-AF_exp_| and the combined standard deviation was calculated using the root of the average measurement variance of the five samples. Muller plots were constructed using ggmuller in RStudio v2023.06.0.

## Data availability

All sequencing reads and hybrid assemblies are available in the National Center for Biotechnology Information (NCBI) under BioProject: PRJNA1288521. List of samples and NCBI accessions are available in **File S3**.

## Acknowledgments

This work was supported by the National Institutes of Health (R21AI166581 to M.C.). We are grateful to Mitra M. Elgrail for assisting with plate sampling.

## Table and Figure captions

**Figure S1.**
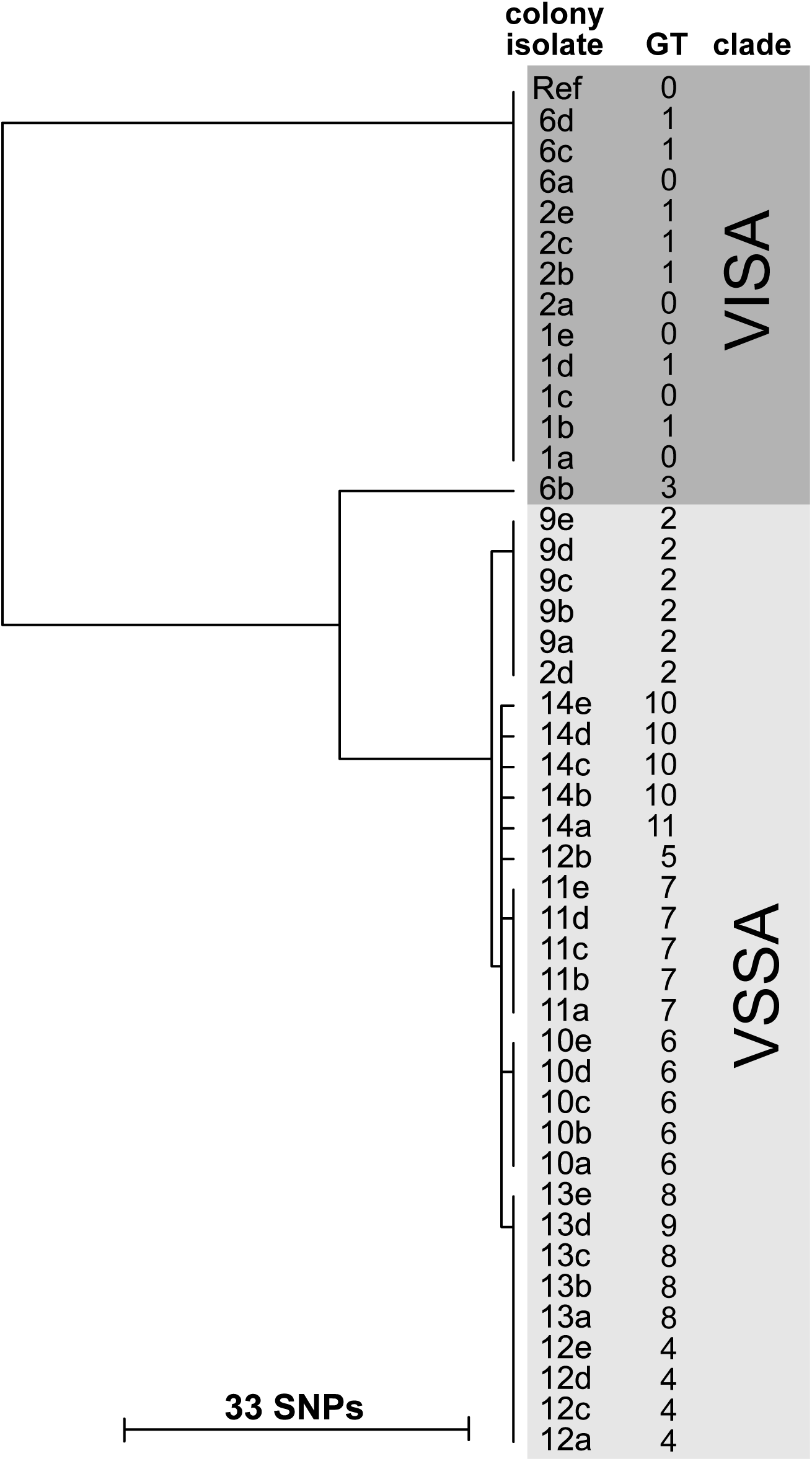
Phylogenetic analysis of Patient B isolates. *snippy* was used to construct the phylogenetic tree. The genotype (GT) of each isolate based on *breseq* variant calls is shown for reference to Figure 6B and the supplementary data files. Isolates grouped into two major clades associated with vancomycin-susceptibility phenotype, indicated by the shaded boxes (VISA, VSSA). Tree scalebar represents the number of single nucleotide polymorphism (SNP) differences.

**Table S1. Patient characteristics.** Abbreviations: history of (h/o), complicated by (c/b), Type 1 diabetes mellitus (T1DM), Type 2 diabetes mellitus (T2DM), end-stage renal disease (ESRD), aortic valve replacement (AVR), mitral valve replacement (MVR), myelodysplastic syndrome (MDS), living donor kidney transplant (LDKT), rheumatoid arthritis (RA), alcohol use disorder (AUD), chronic obstructive pulmonary disease (COPD), renal cell carcinoma (RCC), polysubstance use disorder (PSUD), intravenous drug use (IVDU), native vertebral osteomyelitis (NVO), prosthetic valve endocarditis (PVE), native valve endocarditis (NVE), central line associated bloodstream infection (CLABSI), deep vein thrombosis (DVT), surgical site infection (SSI), central nervous system (CNS), pneumonia (PNA), spinal epidural abscess (SEA); ceftaroline (CPT), daptomycin (DAP), gentamicin (GEN), rifampin (RIF), vancomycin (VAN).

**Table S2. Antibiotic susceptibility tests.** Abbreviations: rifampin (RIF), daptomycin (DAP), ceftaroline (CPT), vancomycin (VAN), zone of inhibition diameter (ZOI), susceptible (S), intermediate (I), resistant (R), non-susceptible (NS).

**File S1. Genotypes and frequency data for Muller plots.**

**File S2. List of variant calls.**

**File S3: Sample Collection with NCBI Accessions.**

## References

1. Culyba MJ, Van Tyne D. 2021. Bacterial evolution during human infection: Adapt and live or adapt and die. PLoS Pathog 17:e1009872.

2. Raghuram V, Gunoskey JJ, Hofstetter KS, Jacko NF, Shumaker MJ, Hu YJ, Read TD, David MZ. 2023. Comparison of genomic diversity between single and pooled Staphylococcus aureus colonies isolated from human colonization cultures. Microb Genom 9.

3. Giulieri SG, Guerillot R, Duchene S, Hachani A, Daniel D, Seemann T, Davis JS, Tong SYC, Young BC, Wilson DJ, Stinear TP, Howden BP. 2022. Niche-specific genome degradation and convergent evolution shaping Staphylococcus aureus adaptation during severe infections. Elife 11.

4. Young BC, Wu CH, Gordon NC, Cole K, Price JR, Liu E, Sheppard AE, Perera S, Charlesworth J, Golubchik T, Iqbal Z, Bowden R, Massey RC, Paul J, Crook DW, Peto TE, Walker AS, Llewelyn MJ, Wyllie DH, Wilson DJ. 2017. Severe infections emerge from commensal bacteria by adaptive evolution. Elife 6.

5. Stoler N, Nekrutenko A. 2021. Sequencing error profiles of Illumina sequencing instruments. NAR Genom Bioinform 3:lqab019.

6. Lieberman TD, Flett KB, Yelin I, Martin TR, McAdam AJ, Priebe GP, Kishony R. 2014. Genetic variation of a bacterial pathogen within individuals with cystic fibrosis provides a record of selective pressures. Nat Genet 46:82–7.

7. Lamy B, Dargere S, Arendrup MC, Parienti JJ, Tattevin P. 2016. How to Optimize the Use of Blood Cultures for the Diagnosis of Bloodstream Infections? A State-of-the Art. Front Microbiol 7:697.

8. Deatherage DE, Barrick JE. 2014. Identification of mutations in laboratory-evolved microbes from next-generation sequencing data using breseq. Methods Mol Biol 1151:165–88.

9. Wood DE, Salzberg SL. 2014. Kraken: ultrafast metagenomic sequence classification using exact alignments. Genome Biol 15:R46.

10. Kim YK, Eom Y, Kim E, Chang E, Bae S, Jung J, Kim MJ, Chong YP, Kim SH, Choi SH, Lee SO, Kim YS. 2023. Molecular Characteristics and Prevalence of Rifampin Resistance in Staphylococcus aureus Isolates from Patients with Bacteremia in South Korea. Antibiotics (Basel) 12.

11. Murphy CK, Mullin S, Osburne MS, van Duzer J, Siedlecki J, Yu X, Kerstein K, Cynamon M, Rothstein DM. 2006. In vitro activity of novel rifamycins against rifamycin-resistant Staphylococcus aureus. Antimicrob Agents Chemother 50:827–34.

12. Ma H, Cheng J, Peng L, Gao Y, Zhang G, Luo Z. 2020. Adjunctive rifampin for the treatment of Staphylococcus aureus bacteremia with deep infections: A meta-analysis. PLoS One 15:e0230383.

13. Baddour LM, Wilson WR, Bayer AS, Fowler VG, Jr., Tleyjeh IM, Rybak MJ, Barsic B, Lockhart PB, Gewitz MH, Levison ME, Bolger AF, Steckelberg JM, Baltimore RS, Fink AM, O’Gara P, Taubert KA, American Heart Association Committee on Rheumatic Fever E, Kawasaki Disease of the Council on Cardiovascular Disease in the Young CoCCCoCS, Anesthesia, Stroke C. 2015. Infective Endocarditis in Adults: Diagnosis, Antimicrobial Therapy, and Management of Complications: A Scientific Statement for Healthcare Professionals From the American Heart Association. Circulation 132:1435–86.

14. Elgrail MM CE, Shaffer MG, Srinivasa V, Griffith MP, Mustapha MM, Shields RK, Van Tyne D, Culyba MJ. 2022. Convergent evolution of antibiotic tolerance in patients with persistent methicillin-resistant *Staphylococcus aureus* bacteremia. Infection and Immunity Accepted Feb 12.

15. Villar M, Marimon JM, Garcia-Arenzana JM, de la Campa AG, Ferrandiz MJ, Perez-Trallero E. 2011. Epidemiological and molecular aspects of rifampicin-resistant Staphylococcus aureus isolated from wounds, blood and respiratory samples. J Antimicrob Chemother 66:997–1000.

16. Camacho C, Coulouris G, Avagyan V, Ma N, Papadopoulos J, Bealer K, Madden TL. 2009. BLAST+: architecture and applications. BMC Bioinformatics 10:421.

17. Hammer ND, Reniere ML, Cassat JE, Zhang Y, Hirsch AO, Indriati Hood M, Skaar EP. 2013. Two heme-dependent terminal oxidases power Staphylococcus aureus organ-specific colonization of the vertebrate host. mBio 4.

18. Ma P, Luo T, Ge L, Chen Z, Wang X, Zhao R, Liao W, Bao L. 2021. Compensatory effects of M. tuberculosis rpoB mutations outside the rifampicin resistance-determining region. Emerg Microbes Infect 10:743–752.

19. Supandy A, Mills EG, Fam KT, Shields RK, Hang HC, Van Tyne D. 2025. Allele-specific effects of mutations in the rifampin resistance-determining region (RRDR) of RpoB on physiology and antibiotic resistance in Enterococcus faecium. mSphere 10:e0050625.

20. Gao W, Cameron DR, Davies JK, Kostoulias X, Stepnell J, Tuck KL, Yeaman MR, Peleg AY, Stinear TP, Howden BP. 2013. The RpoB H(4)(8)(1)Y rifampicin resistance mutation and an active stringent response reduce virulence and increase resistance to innate immune responses in Staphylococcus aureus. J Infect Dis 207:929–39.

21. Nodwell MB, Koch MF, Alte F, Schneider S, Sieber SA. 2014. A subfamily of bacterial ribokinases utilizes a hemithioacetal for pyridoxal phosphate salvage. J Am Chem Soc 136:4992–9.

22. Eren KK, Çınar E, Karakurt HU, Özgür A. 2023. Improving the filtering of false positive single nucleotide variations by combining genomic features with quality metrics. Bioinformatics 39.

23. Cooper VS, Honsa E, Rowe H, Deitrick C, Iverson AR, Whittall JJ, Neville SL, McDevitt CA, Kietzman C, Rosch JW. 2020. Experimental Evolution In Vivo To Identify Selective Pressures during Pneumococcal Colonization. mSystems 5.

24. Lee A, Mirrett S, Reller LB, Weinstein MP. 2007. Detection of bloodstream infections in adults: how many blood cultures are needed? J Clin Microbiol 45:3546–8.

25. Pollitt EJG, Szkuta PT, Burns N, Foster SJ. 2018. Staphylococcus aureus infection dynamics. PLoS Pathog 14:e1007112.

26. McVicker G, Prajsnar TK, Williams A, Wagner NL, Boots M, Renshaw SA, Foster SJ. 2014. Clonal expansion during Staphylococcus aureus infection dynamics reveals the effect of antibiotic intervention. PLoS Pathog 10:e1003959.

27. Prunier AL, Malbruny B, Laurans M, Brouard J, Duhamel JF, Leclercq R. 2003. High rate of macrolide resistance in Staphylococcus aureus strains from patients with cystic fibrosis reveals high proportions of hypermutable strains. J Infect Dis 187:1709–16.

28. Goodwin S, Gurtowski J, Ethe-Sayers S, Deshpande P, Schatz MC, McCombie WR. 2015. Oxford Nanopore sequencing, hybrid error correction, and de novo assembly of a eukaryotic genome. Genome Res 25:1750–6.

29. Wick RR, Judd LM, Gorrie CL, Holt KE. 2017. Unicycler: Resolving bacterial genome assemblies from short and long sequencing reads. PLoS Comput Biol 13:e1005595.

30. Seemann T. 2014. Prokka: rapid prokaryotic genome annotation. Bioinformatics 30:2068–9.

31. Seemann T. 2015. snippy: fast bacterial variant calling from NGS reads. https://github.com/tseemann/snippy. Accessed

32. Letunic I, Bork P. 2024. Interactive Tree of Life (iTOL) v6: recent updates to the phylogenetic tree display and annotation tool. Nucleic Acids Res 52:W78–W82.

33. CLSI. 2024. Performance Standards for Antimicrobial Susceptibility Testing, 34th ed. CLSI supplement M100. Clinical and Laboratory Standards Institute.

